# Adolescent and adult mice use both incremental reinforcement learning and short term memory when learning concurrent stimulus-action associations

**DOI:** 10.1101/2024.04.29.591768

**Authors:** Juliana B. Chase, Liyu Xia, Lung-Hao Tai, Wan Chen Lin, Anne G.E. Collins, Linda Wilbrecht

## Abstract

Computational modeling has revealed that human research participants use both rapid working memory (WM) and incremental reinforcement learning (RL) (RL+WM) to solve a simple instrumental learning task, relying on WM when the number of stimuli is small and supplementing with RL when the number of stimuli exceeds WM capacity. Inspired by this work, we examined which learning systems and strategies are used by adolescent and adult mice when they first acquire a conditional associative learning task. In a version of the human RL+WM task translated for rodents, mice were required to associate odor stimuli (from a set of 2 or 4 odors) with a left or right port to receive reward. Using logistic regression and computational models to analyze the first 200 trials per odor, we determined that mice used both incremental RL and stimulus-insensitive, one-back strategies to solve the task. While these one-back strategies may be a simple form of short-term or working memory, they did not approximate the boost to learning performance that has been observed in human participants using WM in a comparable task. Adolescent and adult mice also showed comparable performance, with no change in learning rate or softmax beta parameters with adolescent development and task experience. However, reliance on a one-back perseverative, win-stay strategy increased with development in males in both odor set sizes. Our findings advance a simple conditional associative learning task and new models to enable the isolation and quantification of reinforcement learning alongside other strategies mice use while learning to associate stimuli with rewards within a single behavioral session. These data and methods can inform and aid comparative study of reinforcement learning across species.

**Author summary:** Here we studied the strategies and mechanisms mice use to learn a simple two choice odor based task in a single session. Using a set size manipulation and computational models we find evidence that mice use incremental reinforcement learning as well as several short-term (one-back) strategies to earn water reward. Our data and models clarify how mice learn a simple task and establish methods by which mouse and human reinforcement learning may be isolated for cross-species comparison of learning.

## Introduction

Mice are the species of choice in many neuroscience experiments due to their genetic tractability, yet questions remain regarding their value as a model for human brain function. As cognitive dysfunction and differences in learning are increasingly implicated in a range of psychiatric and neurodevelopmental disorders, it is essential to understand if mice rely on similar processes as humans to calibrate mouse to human translation. Here, we studied if mice use short-term memory or other strategies alongside reinforcement learning when learning new stimuli-action associations in an odor-based two alternative forced choice (2AFC) task.

There is positive evidence that rodents have working memory, an effortful and flexible form of short-term memory typically associated with higher level executive functions in humans ([1]). Rodents can solve tasks that require correct responses to a cue after a delay lasting dozens of seconds ([2]). For example, in studies of spontaneous alternation and delay-non-match-to-sample, rodents are able to remember where they were seconds to minutes before and change their response ([3]). In these same tasks, there is evidence for increases in rodent working memory in the transition from adolescent to young adult ([4–7]). Despite this evidence, it remains unclear if rodents rely on working memory during simple reinforcement learning tasks where they are rewarded for learning cue-action associations.

It has been noted that in human participants working memory can play a leading role in learning, even in simple tasks which are not intended to tap working memory. Recent evidence has highlighted that much of what is typically considered “reinforcement learning” behavior, where participants learn cue-action associations from rewards, actually relies on working memory processes ([8, 9]). In this work, rapid, yet capacity limited working memory (WM) processes were shown to be used first to solve the task (i.e. when learning about two cues in a block) while incremental but robust reinforcement learning (RL) processes were only additionally revealed when working memory’s limited capacity was exceeded (i.e. when learning more than 3-5 cues in a block, depending on individuals) ([8]). The discovery of contributions of WM to RL was facilitated by the design of a trial-by-trial learning task with set size (the number of stimuli in a block) manipulation and a novel computational model called RL+WM. The task design and model can be used to estimate learning supported by either WM related processes or RL processes in different set sizes ([8]). Interestingly, in a developmental study of human subjects aged 8-30 using the RL+WM task and model, both WM and RL metrics were found to change during adolescent development ([10]), changes likely reflecting the maturation of both the prefrontal cortex and the basal ganglia ([11]).

Here, we adapted the RL+WM task for mice in order to test if 1) working memory is used alongside reinforcement learning in mice during cue-action learning, and whether 2) mice change how they learn the task during adolescent development.

Based on the above data from the human and rodent literature, we predicted that we would see a signature of WM use in mice while learning at low, but not high, set sizes in a rodent version of the RL+WM task. We also hypothesized that the reliance on WM in learning would increase throughout adolescent development, based on past rodent and human work ([4–7, 10]) .

We trained mice in an odor based RL+WM task, in which mice successfully learned new odors in multiple set sizes and within a single session at all ages tested. Following behavioral and regression analyses, we developed a series of models outside of RL+WM to examine unique features of mouse behavior. Based on these models, we concluded mice use basic incremental RL processes along with simple “one-back” strategies at the policy level to solve the task, potentially suggesting a primitive form of short-term memory dependence in learning that falls short of human-like working memory. Some one-back strategies became more commonly used with age in males while the use of others changed significantly over sessions, suggesting a role of sex, development, and experience in shaping short-term strategies. Overall, these data reveal important species differences in neuro-cognitive systems used for learning a simple task and suggest methods for isolating more comparable processes for cross-species comparison.

## Materials and methods

### Animals

C57BL/6 mice were bred and maintained in our colony on a reverse light/dark cycle (12:12 hours) and were group housed throughout experimentation. A total of 47 animals (32 male) were trained on the RL+WM task at ages ranging from P30 to P90 and were water restricted to 85-90% of baseline weight during behavioral studies. All animals were monitored daily for signs of sickness or distress by trained veterinary staff and experimenters. All procedures were approved by the Office of Laboratory and Animal Care at the University of California, Berkeley.

### Behavioral Task

Custom-built operant chambers that have an initiation port in the center and two side ports (see Fig. 1B) were used throughout behavioral experiments. Each port contains an infrared photodiode/phototransistor pair to accurately time port entry and exit (Island Motion, Tappan, NY). Side ports release water through valves (Neptune Research) calibrated to deliver 2 μL of water for correct choices. Slow and constant air flow (0.5L/min) through the center port was redirected through one of several odorant filters (Target2 GMF F2500-19) at trial initiation. Our set-up allows for the delivery of multiple odors independently, creating a seamless transition between learning programs without experimenter intervention.

**Fig 1.**
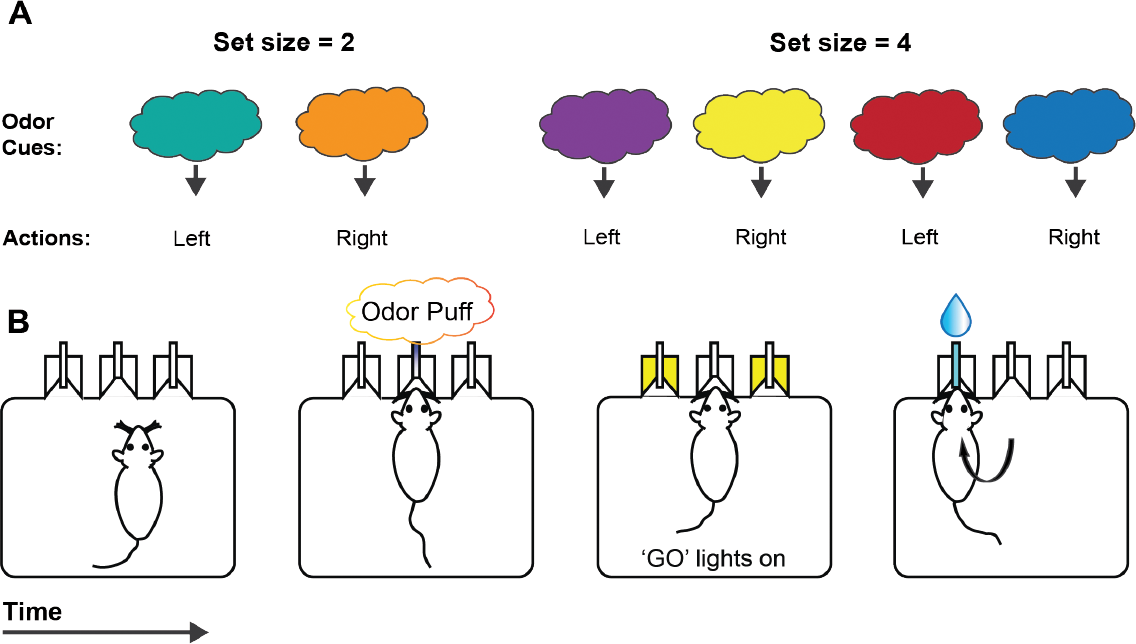
Behavioral schematic of RL+WM task. A) Mice learned to associate an odor cue with a left or right action. After mice pass a readiness criterion (a consistent daily odor pair), they are exposed to novel odors in either set size of 2 odors or 4 odors (presented individually in pseudorandom odor) for that session, that they will only experience in that single session. B) In the operant chamber, mice initiate a trial via nosepoke to the center port where they receive a puff of a single odorant at a time. “GO” lights in the two peripheral ports indicate the availability of water, which mice receive only if they make a correct choice.

**Fig 2.**
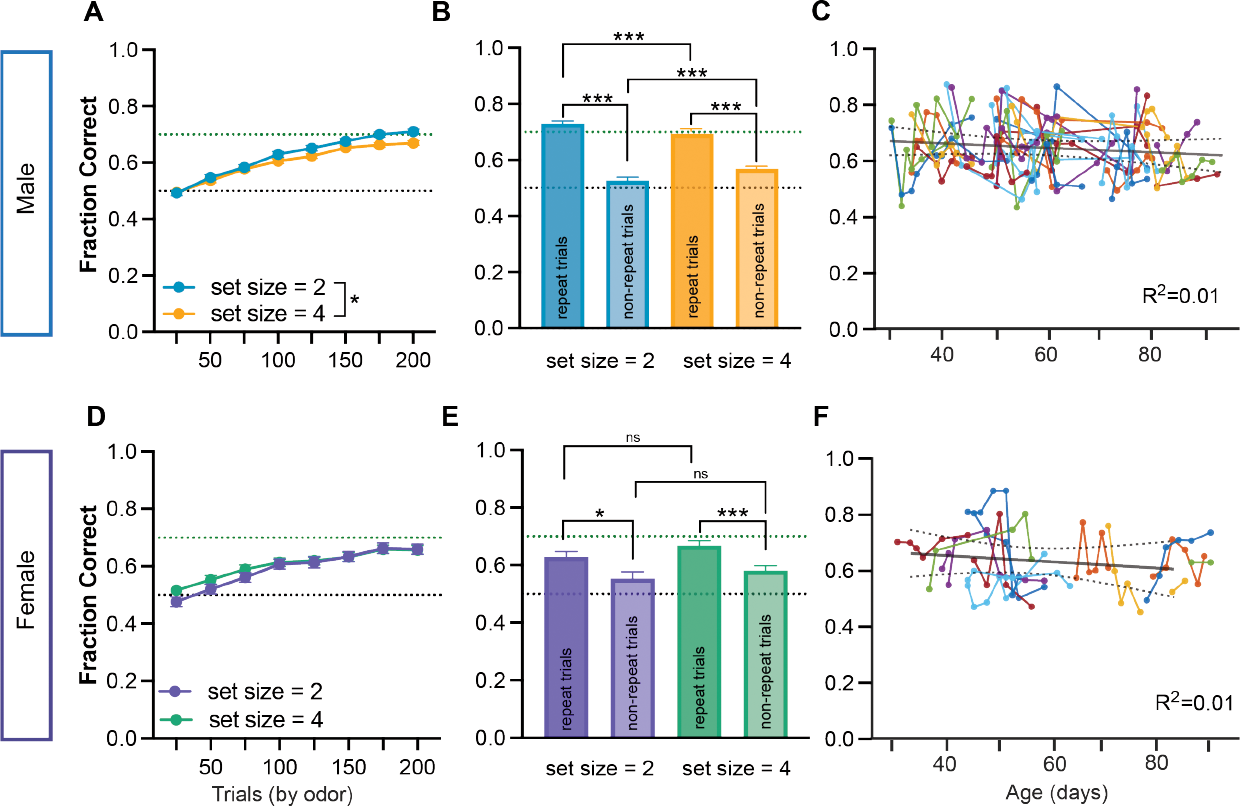
RL+WM data showing from male and female mice learning novel odors across development. Within the first 200 trials per odor, both males (A) and females (D) showed an average performance (fraction correct) significantly above chance (dotted black line at 50%, dotted green line at 70%) in both set size = 2 and set size = 4. Males showed a small but significant difference in performance between set sizes driven by the 150-200th presentation of each novel odor. Female mice had no significant difference between set size = 2 and set size = 4. (B, E) Performance in males and females showed significant benefit when an odor stimulus was repeated two trials in a row when compared to non-repeated stimulus trials. This effect was observed in both set sizes for both sexes. However, in male mice, while performance in repeat trials was higher in set size = 2 than in set size = 4 (possibly driven by a higher proportion of repeat trials), this relationship was reversed when performance in non-repeat trials was examined. (C,F) Fraction correct in males and females did not change across development P30-90, a period that includes both adolescent development and early young adulthood in mice. Lines connect up to three sessions per mouse and show performance for first 200 trials per odor in both set size = 2 and set size = 4 for all sessions analyzed. Dotted lines indicate SEM. Asterisks indicate the result of RM 2-way ANOVA (A,D) and Wilcoxon signed rank test (B,E). *** *p <* 0.001, ** *p <* 0.01, * *p <* 0.05.

Inexperienced, water-restricted male and female mice were first pre-trained to initiate trials via nose poke to the center port and to associate side ports with water reward for several days. Next, animals encountered a preliminary odor pair, A(cinnamon) - B(vanilla), which stably predicted the left and right port respectively for reward. Animals were overtrained on A/B and experienced this odor pair at the beginning of each novel odor learning session as a “performance readiness check.” After two or three sessions in this early learning phase mice were then run for at least three sessions with novel odors presented in set sizes of n=2 or n=4 (six sessions total per mouse) in the full RL+WM task (novel odor phase). Following early learning mice were required to reach performance levels of 70% on both sides on A/B (indicating task knowledge) before exposure to novel odors. Each full task session, one per day, included 150-300 presentations of odors A&B followed by 200-400 presentations (dependent on set size) of each novel odor in a pseudo-random manner, controlling for a normal delay distribution between two successive presentations of the same stimulus.

### Behavioral analysis

Animal operant performance for novel odors was analyzed in MATLAB (Mathworks, Natick, MA) using custom scripts and GraphPad Prism. Three sequential sessions for both set size = 2 and set size = 4 were taken from each animal and used for learning curves and behavioral analyses as well as all subsequent analyses and modeling.

### Behavioral statistics

In order to understand how mice learn to associate odor with rewarding choices at a high level, we first performed a series of behavioral analyses on the data using MATLAB (version 2018, 2022b) and GraphPad Prism (version 9.1.1). Processed data with performance information was uploaded in GraphPad Prism, and normality and variance assumptions were checked using Anderson-Darling, D’Agostino and Pearson test, Shapiro-Wilk test, and Welsh’s test. In order to account for variable session length (and allow for comparisons between set sizes), we analyzed the first 200 presentations of each novel odor stimuli of the first 3 sessions of each set size for each animal. Corrections for multiple comparisons were applied used Sidak’s method to control the overall Type 1 error rate. We used the following statistical tests to examine behavioral differences between set sizes, sexes, types of trials, and across development (postnatal day, age, of animal): repeated-measures two-way analysis of variance (RM 2-way ANOVA), post-hoc Sidak’s multiple comparisons tests, Wilcoxon signed rank test, t-tests, and simple linear regression. Significance level was set to *p <* 0.05 for all statistical tests, and confidence intervals as well as effect sizes were reported where applicable. To assess the impact of age on performance, separate mixed linear models were fitted for each sex using statsmodels package in Python. The models included age as the fixed effect predictor variable with additional fixed effects for session number and set size. Random effects (intercept and slope) were included for individual animal identity.

### Logistic regression analysis

To better characterize trial-by-trial choice dynamics, we used logistic regression analysis. We developed multiple models to capture different hypotheses. In all cases, we sought to predict *P*_*t*_(*Correct*) via a logistic link function for each individual session.

In a first regression model, we included predictors that captured theoretical markers of incremental RL, as well as markers of strategies relying on recent information, possibly dependent on WM. The “*correct history*” regressor was meant to capture contributions of an incremental RL process and defined as the number of past correct (reinforced) choices for the current trial’s odor. For WM, we examined the effect of repeating stimulus in recent past trial (1-back = 1 if odor on trial t, *o*_*t*_, is the same as trial t-1, *o*_*t−*1_; 0 otherwise) or two trials back (2-back = 1 if odor on trial t, *o*_*t*_, is the same as trial t-2, *o*_*t−*2_; 0 otherwise).

In further regression models, we considered not only the reward history, but more specifically the characteristics of the last trial. In particular, we categorized all trials as a function of two orthogonal characteristics: stimulus repeating (1-back = 1) vs. non-repeating (1-back = 0), and previously rewarded vs. non-rewarded outcomes.

For example, in regression 3, we additionally modeled interactions between these variables using the following regressors:

- correct history, characterizing incremental learning.
- reward at *t−* 1, characterizing potential immediate performance improvement after a rewarded trial.
- reward-repeat (regressor = 1 if *o*_*t−*1_ = *o*_*t*_ and *r*_*t−*1_ = 1, *−*1 if *o*_*t−*1_ ≠ *o*_*t*_ and *r*_*t−*1_ = 1, 0 if *r*_*t−*1_ = 0), characterizing a WM-like process, whereby having just observed relevant information (correct choice for the current stimulus) improves performance.
- noreward-repeat (regressor = 1 if *o*_*t−*1_ = *o*_*t*_ and *r*_*t−*1_ = 0, *−*1 if *o*_*t−*1_ ≠ *o*_*t and*_ *r*_*t−*1_ = 0, 0 if *r*_*t−*1_ = 1), characterizing a baseline for reward-repeat.

All parameters for within individuals and sessions logistic regressions were estimated using Matlab’s glmfit function or statsmodels package in Python. Following normality tests on individual session regression coefficients, either parametric (one sample t) or nonparametric (Wilcoxson signed rank) tests were run in GraphPad Prism and reported in the text.

### Mixed linear model analysis

To test the influence of development on trial-by-trial learning we determined the relationship between the age and each regression coefficient for males, females, and across both set sizes, using mixed linear models. First, we visualized and standardized the data points by taking z-scores for each session’s coefficient and dropping any outliers, namely animal sessions that were over 3 standard deviations from the mean (7 out of 96 observations). Next, to coarsely test the existence of a relationship between each regression coefficient and age across sessions, we ran a simple linear regression looking at goodness of fit (determined by *R*^2^ and corresponding *p*-value as shown in Fig. 4,5) without controlling for non-independence when multiple measures were taken from a single individual. To assess the normality of each regression coefficient distribution, we employed both quantitative analysis via an Anderson-Darling normality test and qualitative examination through density and Q-Q plots. Following this evaluation, we used either a Kruskal-Wallis test or a one-way ANOVA to investigate the potential influence of animal age on the variance observed in the dependent variable (regression coefficient). After checking mixed linear model common assumptions, such as linearity and the existence of both normally distributed and independent errors, we assessed multicollinearity by calculating the Variance Inflation Factor (VIF) for the predictor variables. The VIF values were examined to ensure that there was no problematic multicollinearity in the model. The statsmodels package in Python 3 was utilized for this analysis. Subsequently, we fitted mixed linear models, incorporating random intercepts and random slopes for each animal to control for repeated measures from each individual. Results from the mixed linear model analysis, including checks for multicollinearity, are reported in the text and/or figure legends.

**Fig 3.**
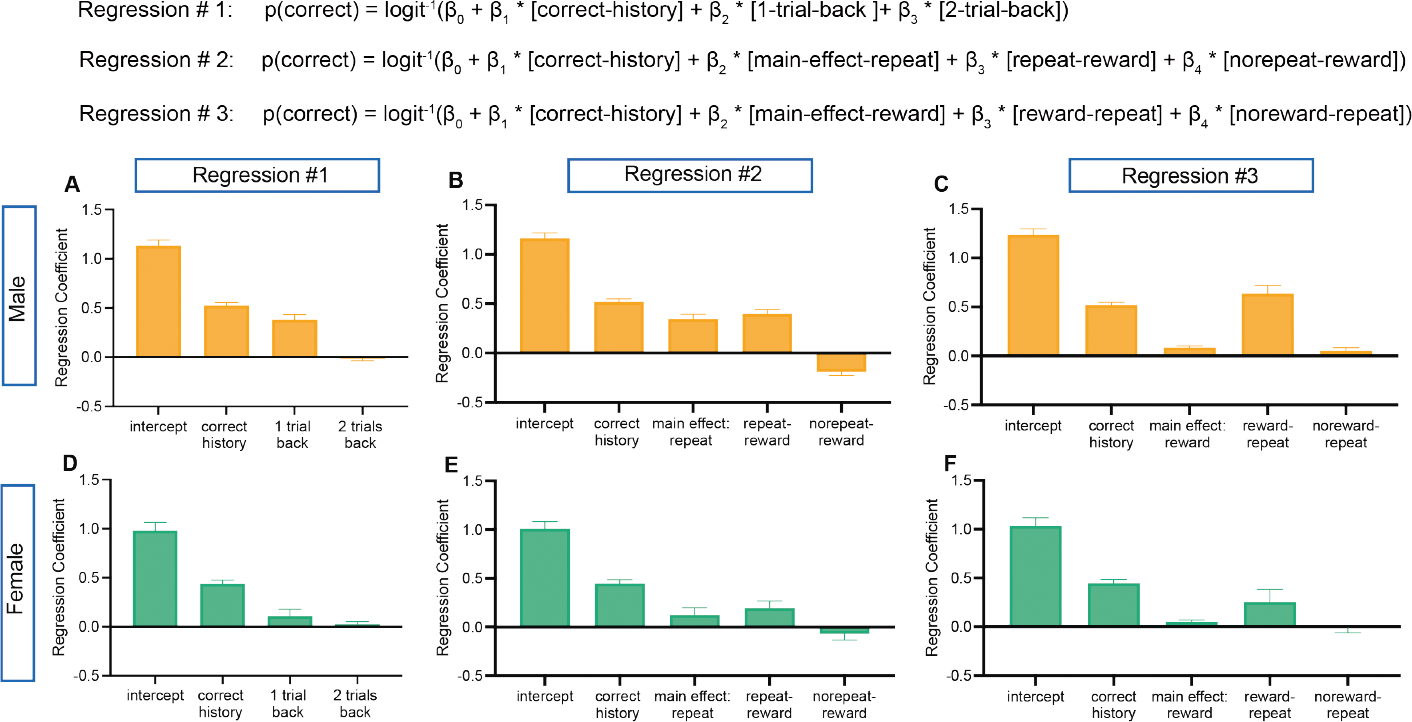
Summary regression coefficients from regressions run in set size = 2. In order to understand the influence of past trials on current trial *t*, three regressions with a logit link function were developed and fit to individual sessions. All regressions included a theoretical marker for RL (correct history) and a series of different one-back trial identities that could be used to approximate WM. Regression analysis with interaction of past trial (A,D) the influence of stimulus repetition (B,E) and the influence of past trial’s reward (C,F) on the current trial are derived and summarized from set size = 2 individual sessions for both male (A-C) and female (D-F) mice. The influence of correct history for both male and female mice and for all three regressions was significant indicating the use of RL. For males, regression #1 (A) *t* = 17.9, *df* = 95, *p <* 0.0001, regression #2 (B) *t* = 18.29, *df* = 95, *p <* 0.0001, regression #3 (C) *t* = 18.29, *df* = 95, *p <* 0.0001. For females, regression #1 (D) *t* = 12.55, *df* = 38, *p <* 0.0001, regression #2 (E) *t* = 12.78, *df* = 38, *p <* 0.0001, regression #3 (F) *t* = 12.78, *df* = 38, *p <* 0.0001. By looking at one-back and two-back trials in regression #1, we found that one-back was significantly above 0 only for male mice and that two-back was not significant for either sex. For males (A) intercept:*t* = 18.8, *df* = 95, *p <* 0.0001; one-back: *W* = 3102, *p <* 0.0001; two-back: *W* =*−* 502, *p* = 0.36. For females (D) intercept: *t* = 11.96, *df* = 8, *p <* 0.0001; one-back: *W* = 140, *p* = 0.33; two-back: *W* = 8, *p* = 0.96. The influence of one-back trials for male mice led us to separate trials into the influence of stimulus (regression #2) and reward (regression #3). For regression #2 (B) for males, there was a significant effect of repeating trials and the interaction between repeating rewarded trials and non-repeating rewarded trials: intercept: *t* = 20.7, *df* = 95, *p <* 0.0001 ; main effect of repeat: *W* = 2938, *p <* 0.0001; repeat - reward: *t* = 8.79, *df* = 95, *p <* 0.0001; no repeat - reward: *W* = *−* 2264, *p <* 0.0001. For regression #2 (E) for females, there was only a significant effect of repeating trials when the previous trial was rewarded: intercept: *t* = 13.69, *df* = 38, *p <* 0.0001; main effect of repeat: *W* = 148, *p* = 0.30; repeat - reward: *t* = 2.5, *df* = 38, *p* = 0.01; no repeat - reward: *W* = *−* 70, *p* = 0.63. For regression #3 (C) for males, all coefficients were significantly above 0: intercept: *t* = 20.57, *df* = 95, *p <* 0.0001; main effect of reward: *t* = 5.52, *df* = 95, *p <* 0.0001, reward - repeat: *W* = 3364, *p <* 0.0001, no reward - repeat: *W* = 1230, *p* = 0.02. For regression #3 for females (F) there was only a significant effect of reward on previous trial: intercept: *t* = 13.06, *df* = 38, *p <* 0.0001; main effect of reward: *t* = 2.10, *df* = 38, *p* = 0.04, reward - repeat: *W* = 148, *p* = 0.30, no reward - repeat, t-test: *t* = 0.03, *df* = 38, *p* = 0.97. Normality tests on grouped regression coefficients from individual sessions were run as described in Materials and methods and parametric (one sample *t*) or non-parametric (Wilcoxon signed rank) tests were applied accordingly. See Figure S3 for corresponding summary for set size = 4.

**Fig 4.**
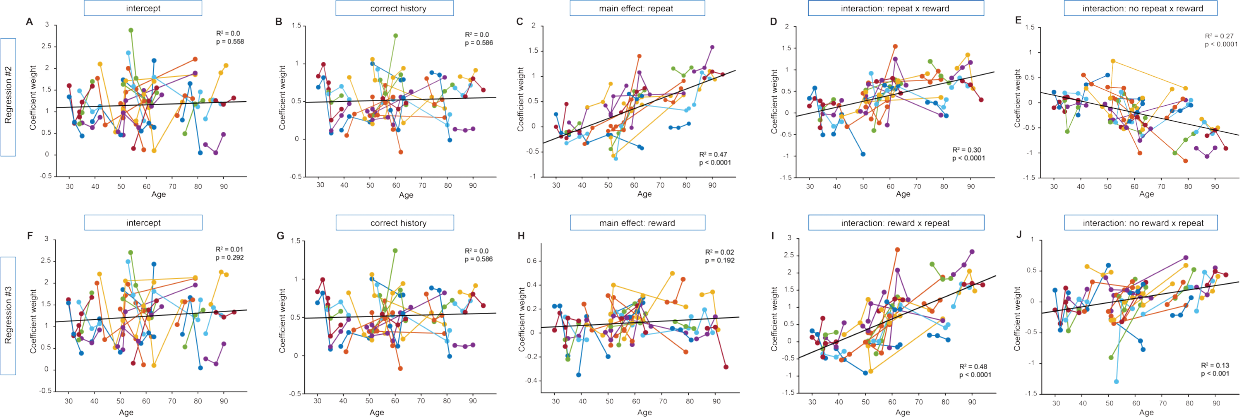
Male regression coefficients across development in set size = 2. In order to understand the relationship between age and the predictors of current choice in trial *t* (see Fig. 3 for the logistic regression summary), we looked at whether coefficient weight (y-axis) would change over development (x-axis) for male mice. It is possible that as mice age, they rely to differing extents on separable learning systems (for example, RL, captured by ‘correct history’ and, WM, captured by one-back combinations of stimuli and reward). For each coefficient from regression #2 (A-E) and regression #3 (F-J), we report the *R*^2^ and *p*-value from a simple linear regression (best fit line) and the results of a mixed linear regression model. This technique allowed us to account for variability caused by repeat (3) sessions of individual mice (each mouse colored separately and individual sessions connected) (see Mixed linear model analysis for more information about analyses). While ‘correct history,’ a measurement of RL, was significantly above 0 for both regressions (Fig. 3), the coefficient weight did not change across age. (B) repeat correct history: *β*_*age*_ = 0.002, 95% CI = [*−*0.002, 0.005], *p* = 0.34 and (G) reward correct history: *β*_*age*_ = 0.002, 95% CI = [*−* 0.002, 0.005], *p* = 0.34. For regression #2, a regression focusing on repeating or non-repeating stimuli, both the coefficient for repeat and repeat - reward increased over development, while no repeat - reward decreased: (A) repeat intercept: *β*_*age*_ = 0.005, 95% CI = [*−*0.002, 0.01], *p* = 0.15; (C) main effect of repeat: *β*_*age*_ = 0.024, 95 % CI = [0.016, 0.033], *p <* 0.0001; (D) repeat - reward: = 0.014, 95 % CI = [0.009, 0.018], *p <* 0.0001; (E) no repeat - reward: *β*_*age*_ = *−* 0.010, 95 % CI = [*−* 0.014, *−* 0.006], *p <* 0.0001. For regression #3, a regression focusing on rewarded or unrewarded trials, both the interaction of repeating stimuli with rewarded or unrewarded previous trials increased across development: (F) reward intercept: *β*_*age*_ = 0.006, 95% CI = [*−*0.001, 0.013], *p* = 0.074; (H) reward main effect: *β*_*age*_ = 0.001, 95% CI = [*−*0.001, 0.003], *p* = 0.28; (I) reward - repeat: *β*_*age*_ = 0.03, 95% CI = [0.022, 0.038], *p <* 0.0001; (J) no reward - repeat: *β*_*age*_ = 0.009, 95% CI = [0.004, 0.013], *p <* 0.0001. To see changes across development in set size = 4, see Fig. S4.

**Fig 5.**
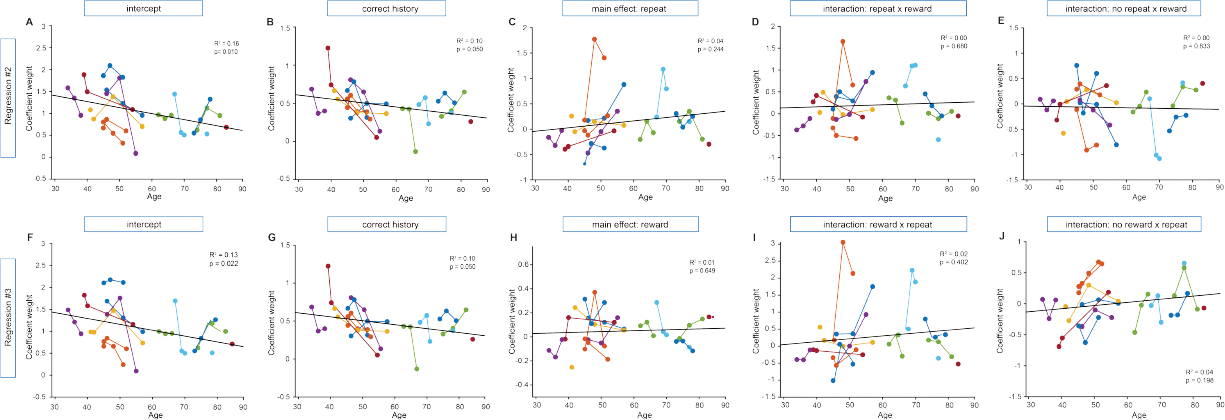
Female regression coefficients for set size = 2 across development. In order to understand the relationship between age and the predictors of current choice in trial *t* (see Fig. 3 for the logistic regression summary), we looked at whether coefficient weight (y-axis) would change over development (x-axis) for female mice and if this would differ from male mice (Fig. 4). It is possible that as mice age, they rely to differing extents on separable learning systems (for example, RL, captured by ‘correct history’ and, WM, captured by one-back combinations of stimuli and reward). For each coefficient from regression #2 (A-E) and regression #3 (F-J), we report the *R*^2^ and *p*-value from a simple linear regression (best fit line) and the results of a mixed linear regression model. This technique allowed us to account for variability caused by repeat (3) sessions of individual mice (each mouse colored separately and individual sessions connected) (see Mixed linear model analysis for more information about analyses). While ‘correct history,’ a measurement of RL, was significantly above 0 for both regressions (Fig. 3), the coefficient weight did not change across age. (B) repeat correct history: *β*_*age*_ = *−*0.004, 95% CI = [*−*0.01, 0.002], *p* = 0.20 and (G) reward correct history: *β*_*age*_ = *−* 0.004, 95% CI = [*−* 0.01, 0.002], *p* = 0.20. There was a slight decrease in the regression coefficient weight of intercept for both regression #2 and regression #3: (A) intercept: *β*_*age*_ = *−*0.014, 95% CI = [*−*0.02, *−*0.003], *p* = 0.017 and (F) intercept: *β*_*age*_ =*−* 0.015, 95% CI = [*−* 0.029, *−* 0.001], *p* = 0.031. Across regression #2, a regression that focused on repeating or non-repeating stimuli, there were no significant changes across development: (C) main effect of repeat: *β*_*age*_ = 0.015, 95% CI = [*−*0.008, 0.039], *p* = 0.20, (D) repeat - reward: *β*_*age*_ = 0.002, 95% CI = [*−*0.01, 0.014], *p* = 0.78, (E) no repeat - reward; *β*_*age*_ = *−* 0.001, 95% CI = [*−* 0.013, 0.011], *p* = 0.86. For regression #3, a regression that focused on rewarded or non-rewarded trials, there were no significant changes across development: (H) main effect of reward: *β*_*age*_ = 0.00, 95% CI = [*−*0.004, 0.005], *p* = 0.82, (I) reward - repeat: *β*_*age*_ = 0.01, 95% CI = [0.01, 0.032], *p* = 0.344, (J) no reward - repeat: *β*_*age*_ = 0.007, 95% CI = [*−*0.003, 0.017], *p* = 0.145. These results are in contrast to significant changes seen across development in male mice (Fig. 4), but mimic lack of changes observed in female mice in set size = 4 (S4).

### Computational modeling

#### Model specifications

To investigate the processes that support the animals’ learning and decision making process, we used computational modeling. All models investigated included a reinforcement learning (RL) component, under the assumption that animals used reward outcomes *r* = 0, 1 to learn to estimate the value *Q*(*o, a*) of making choice *a* (left/right) for odor *o*, using a standard delta rule:

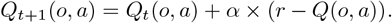

Values were initialized to be uninformative at *Q*_0_ = 0.5. Choice was determined through a softmax policy over estimated values: *P* (*a*|*o*) *∝* exp (*β*(*Q*(*o, a*))), where inverse temperature *β* indexed the degree of stochasticity in choosing the best option.

Beyond this baseline model, we considered whether computational models with additional mechanisms could better account for the data. Mechanisms we considered included learning rate asymmetry, forgetting, and mixture of RL with 1-back strategies. We detail these mechanisms below. Note that we did not include the WM mechanisms considered in human RL-WM models, because the very limited set size effects observed in mice indicate that human-like WM mechanisms are not at play here.

#### Learning rate asymmetry

is frequently considered in RL models and often improves fit. Here, we considered two possibilities: setting *α*_+_ and *α*_*−*_ as two free parameters applied to positive and negative prediction errors, respectively. Based on the results of this model where we found very low values of *α*_*−*_, we also considered a family of models with fixed *α*_*−*_ = 0.

#### Forgetting

is implemented as a decay of all odor-action values towards initial values at every trial: *Q*_*t*+1_ *← Q*_*t*_ + *ϕ ×* (*Q*_0_ *− Q*_*t*_), with decay parameter 0 *≤ ϕ ≤* 1.

#### One-back strategies

Previous work has shown that simple changes in the policy of RL models may often strongly improve fit to behavior. The simplest such strategy is a “choice kernel”, or “sticky choice”, whereby the mice tend to repeat the previously chosen action, independently of stimulus and outcome. This is implemented by modifying the policy such that

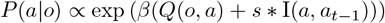

where I(*a, a*_*t−*1_)) = 1 if *a* = *a*_*t−*1_, 0 otherwise, and *s* is a free parameter capturing degree of tendency to repeat (or avoid) the previous trial’s choice.

We additionally considered slightly more complex one-back strategies that incorporated information about the last trials’ stimulus and outcome in addition to previous choice, capturing a rudimentary form of working memory. Specifically, we considered four types of trials *T*_*i*_; *i* = 1 : 4, as a function of whether the previous trial was the same or a different odor and whether the previous trial was rewarded or unrewarded. Trial-type dependent strategies were expressed as a bias on the policy following:

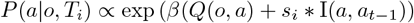

Fig. 6A describes the 4 types of trials and corresponding policies. We considered various models with subsets of those policies set to 0 or to each other’s values (e.g. *s*_2_ = *s*_4_).

**Fig 6.**
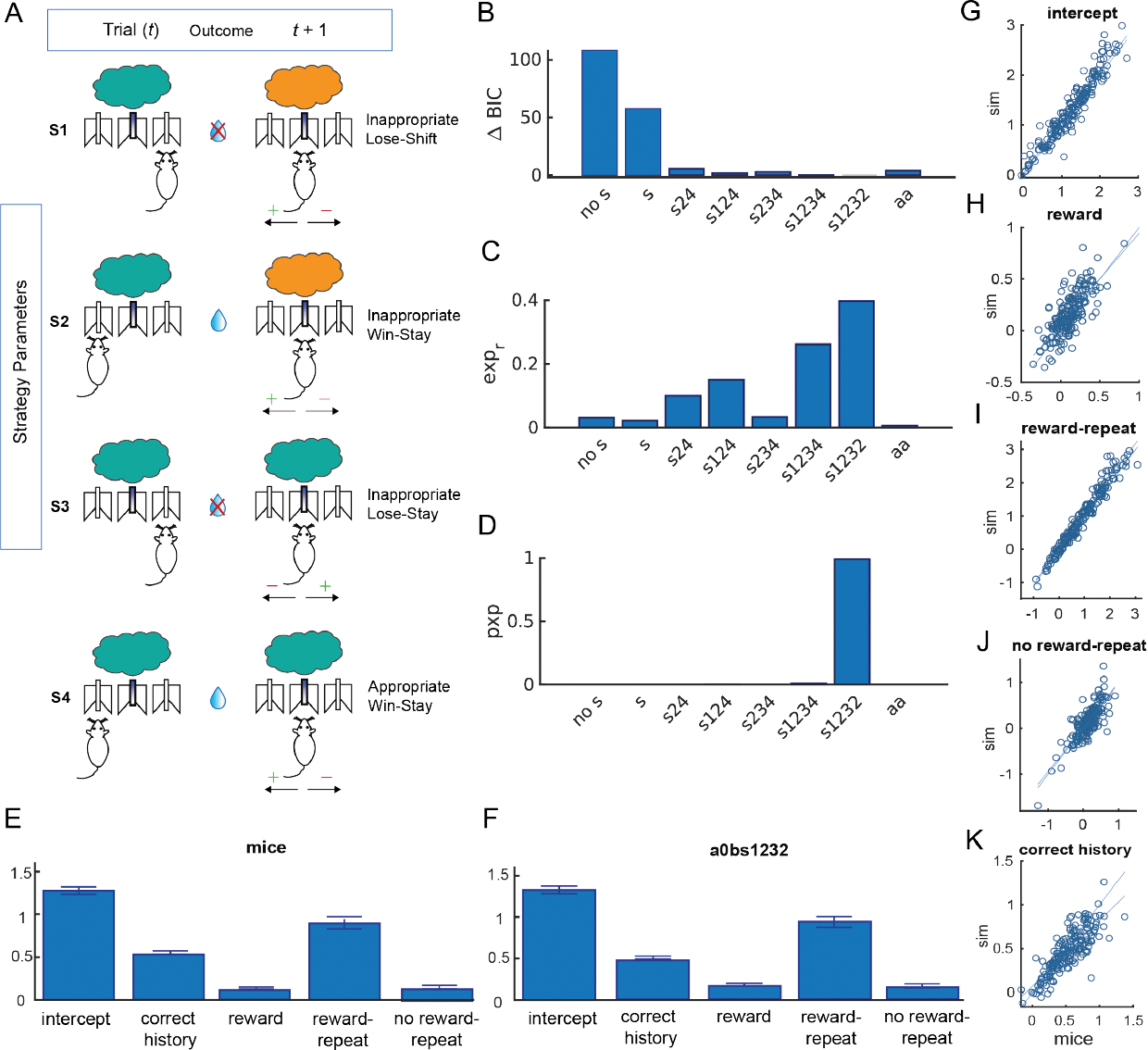
Model Comparison and Validation. (A) We examined 4 parameters that isolated different ways mice might use 1-trial back information (S1-S4). Later in the best-fitting model, S2 and S4 were collapsed to one parameter to reduce complexity. Each left cartoon indicates the stimulus and the mouse’s action at time *t* and the cartoon on the right reflects the choice options at time *t* +1 that a mouse has when presented with either stimulus. The plus/minus sign reflect the positive/negative direction of the specific strategy parameter. The descriptions of the strategy parameters describe the positive value of the parameter at time *t* +1. (B-D) The winning model a0bs1232 (where a indicates a free *α*_+_ parameter, 0 indicates that *α*_*−*_ = 0 is fixed, and s1232 indicates three free S parameters with S4=S2) was compared to other a0b models with various combinations of strategy parameters (s) or no strategy parameters (nos) as well as a model leaving the negative learning rate free (aabs1232). All competing models built from basic RL model described in Materials and methods. The winning model had an average BIC of 859.4, not significantly different from the full model a0bs1234 (BIC=857.6191). However, a Bayesian model comparison analysis showed that a0bs1232 was more frequent across sessions (39.74% vs. 26.11% for a0bs1234), with a protected exceedance probability of 0.991, confirming that it was the best model at the group level. (E-F) Example logistic regression. To validate our winning model we analyzed simulated behavior from parameters fit on each individual and session with the same logistic regression approach. We found that the simulated behavior captured well the impact of repeat, reward, their interaction and the effect of reward history on a session by session basis, as shown by the group level effect (E-F) and tight correlations between mouse behavior and simulated behavior using fit parameters (sim; G-K).

Our winning model included 5 free parameters: *β, α*_+_, *s*_1_,*s*_2_,*s*_3_, as well as two fixed parameters (*α*_*−*_ = 0 and *s*_4_ = *s*_2_).

#### Model fitting

We used standard maximum likelihood estimation to estimate best-fitting parameters [12]. While we considered using more state-of-the art methods such as *maximum a posteriori* or hierarchical Bayesian modeling, we found this unnecessary in this case, as the large number of trials afforded very good parameter identification (tested through a generate and recover procedure) with the simpler method, with no need for the additional assumptions that more advanced methods require (e.g. priors). We fit separately each individual and session, obtaining a set of best fit parameters for each individual and session.

### Model comparison

We computed the Bayesian Information Criterion (BIC, [13]), and also performed Bayesian model selection via the protected exceedance probability ([14]), using SPM12’s `spm_bmf` Matlab function (Fig. 6B-D). Bayesian model selection results in a more robust model comparison than other fixed effects comparison strategies. Confusion matrices ([12]) confirmed that this process supported good model identifiability.

### Model validation

To confirm that our model was able to capture mouse behavior adequately, we simulated the winning model with fit parameters (see Fig. S1), and performed similar analyses as on empirical data. For example, we were able to capture well not only group level learning curves (see Fig. S2), but also trial-by-trial dynamics at the individual and session level, as measured by the logistic regression analysis (Fig. 6E-F) .

### Mixed linear model analysis of model parameters

To determine the relationship between age and parameter weight for each sex and set size, mixed linear models were used as explained above(Mixed linear model analysis). Both *α*_+_ and *β* were log-transformed to log(*α*_+_) and log(*β*) prior to analysis to achieve a normal distribution and resolve issues of non-constant variance.

## Results

### Behavioral task and results

Mice were trained in the RL+WM task using novel odors at ages ranging from P30 to P100. Pre-training to perform the task was established in all mice before novel odors were presented. Water-restricted mice were first habituated to the chamber, the central task initiation port and its two peripheral water delivery ports (Fig. 1B). Once nose poking and drinking behavior were established, mice entered the early training phase where they were trained with the first set of odors “A & B,” delivered at the central initiation port with a left choice rewarded for odor A and right choice rewarded for odor B (Fig. 1A). After two to three sessions in the early training phase, mice were then run for at least three sessions with novel odors presented in set sizes of either n=2 or n=4 (six sessions total per mouse) in the RL+WM task (novel odor phase) (Fig. 1A). Performance “readiness” was checked at the beginning of each session by presentation of 150-300 odor A & B trials. To move on to novel odors, mice had to reach a 70% performance criterion for odor A & B. If they failed to reach this performance readiness criterion they were only exposed to odor A & B for that session.

In the novel odor learning phase of RL+WM, we found evidence of significant learning within a single session in both males and females of all ages. For each new odor, mice started near chance performance, but over the course of two hundred trials per odor they approached an average of 70% performance (Fig. 2A,D).

In humans performing RL+WM, stimuli encountered in lower set sizes (e.g. two cues) have been shown to be learned quickly using WM, while stimuli in higher set sizes (e.g. 6 cues) are learned more gradually with greater reliance on RL systems. In both male and female mice (all ages combined), we found that learning was similarly gradual for odors presented in set sizes of two or four odors (Fig. 2A,B). Male mice had a small but significant difference in performance between set size 2 and set size 4 that emerged approximately following the 150th presentation of each odor (Fig. 2A, RM 2-way ANOVA, set size: *F* (1, 520) = 22.58, *p <* 0.0001, Holm-S?idáak: 175th trial per odor, *p* = 0.02, 200th bin, *p* = 0.01, Fig. 2A). The weak set size effect in males was reversed when considering only non-repeat trials (by performance, Mann-Whitney *U* = 7507, *p <* 0.0001, Fig. 2B), indicating that the slightly better overall learning in set size 2 compared to set size 4 is likely driven by the higher proportion of repeat trials in set size 2. Performance in set size = 2 was not significantly different from performance in set size = 4 in female mice (*t*-test: *t* = 0.21, *df* = 93, *p* = 0.83, Fig. 2D) and there was no significant difference between non-repeat trials (*t* -test: *t* = 0.88, *df* = 100, *p* = 0.37, Fig. 2D).

Males and females had similar behavioral performance in the task (Mann-Whitney, *U* = 14046, *p* = 0.19), both showing increased performance in repeat versus non-repeated trials in both set sizes (Wilcoxon signed rank test: male: *W* =*−* 18971, *p <* .0001 for set size = 2 & *W* = *−* 993, *p <* .0001 for set size = 4; female: *W* = *−* 630, *p* = 0.017 for set size = 2 & *W* = *−* 606, *p <* .001 for set size = 4, Fig. 2B, E.).

Performance on the task was also comparable across all adolescent and adult ages tested (mixed-effect linear regression, male: *β*_*age*_ = *−*0.001, 95 % CI = [*−* 0.002, 0], *p* = 0.207; female: *β*_*age*_ = *−* 0.001, 95 % CI = [*−* 0.003, 0.002], *p* = 0.514, Fig. 2C, F).

Based on the minimal effect of set size in mice, which was mostly driven by repeat trials, we inferred that learning in both set sizes was likely mostly supported by the RL learning system, as a human-like efficient working-memory component would have elicited strong set size effects, including in non-repeat trials. Furthermore, unlike the emergence of a set size difference following the 150th presentation of each stimuli that we observed in mice, human-like working memory would be expected to contribute most to performance early in learning.

### Logistic regression models

To better characterize the effect of recent history separately from the effect of long-term cumulative reinforcement, we next analyzed trial-by-trial choices for each individual session in a basic logistic regression with correct history, 1-back repeated cue, and 2-back repeated cue as predictors (see Materials and methods). We found for set size =2 that there was a significant positive effect of correct history across sessions for both males and females (t-test, males: *t* = 17.9, *df* = 95, *p <* 0.0001, females: *t* = 12.55, *df* = 38, *p <* 0.0001, Fig. 3A, D), confirming that mice were more likely to make a correct choice the more they experienced reward, as expected from an incremental RL process. The regression coefficient for 1-back repeated cue was also significantly above 0 for males, but not for females (Wilcoxon signed rank test, males: *W* = 3102, *p <* 0.0001, females: *W* = 140, *p* = 0.33), but there was no 2-trial back effect for either sex (Wilcoxon signed rank test, males: *W* =*−* 502, *p* = 0.36, females: *W* = 8, *p* = 0.96). This indicates that mice might be maintaining and leveraging information in the immediately previous trial, *t −* 1, but not further in the past. Notably, male animals exhibited a significantly higher one-trial back regression coefficient compared to females (Mann-Whitney test, *U* = 1316, *p* = 0.006), prompting us to continue to analyze male and female data separately in subsequent analyses. Importantly, all regressors that significantly captured animal’s behavior in set size = 2, were also significant in set size = 4 (see Fig. S3A, D). This replicates our behavioral finding that animals in the RL+WM task perform similarly in set size = 2 and set size = 4 sessions.

To further probe which information in the previous trial was used, we categorized all trials as a function of two orthogonal characteristics: repeating cue (1-back = 1) vs. non-repeating cues (1-back = 0), and previously rewarded (reward = 1) vs. non-rewarded (reward = 0) trials. We ran two additional regressions to triangulate the potential interaction between repeated cues and reward accordingly.

In regression model 2 for set size = 2, where we looked at correct history, repeat, and previous reward for repeat and non-repeat trials separately, we confirmed a significant effect of correct history for both sexes (t-test males: *t* = 18.29, *df* = 95, *p <* 0.0001, females: *t* = 12.78, *df* = 38, *p <* 0.0001), supporting RL contributions. We also found a significant effect of repeated cues in males only (Wilcoxon signed rank test, males: *W* = 2938, *p <* 0.0001, females: *W* = 148, *p* = 0.30), supporting male animals’ use of short-term memory of the last trial. Furthermore, there was a significant positive effect of reward in repeat trials for both sexes (t-test, males: *t* = 8.79, *df* = 95, *p <* 0.0001, females: *t* = 2.5, *df* = 38, *p* = 0.01, Fig. 3B,E), as expected if mice can maintain and leverage the correct information observed in the last trial. However, there was also a significant negative effect of reward in non-repeated cue trials for male animals (Wilcoxon signed rank test, males: *W* = *−* 2264, *p <* 0.0001, females: *W* = *−* 70, *p* = 0.63, Fig. 3B,E). The positive effect of reward for repeat trials and negative coefficient for non-repeat ones could suggest, rather than adaptive use of WM or simpler one-back memory, that male mice might be sticking to their choice in trial *t−* 1 if they were rewarded. This would help their performance in trial *t* if it is a repeated cue, but hurt performance if it is non-repeating (a different cue). If this was the case, we would expect the magnitude of the two coefficients to be indistinguishable. However, a paired t-test between the “repeat-reward” and “no-repeat-reward” coefficients for male animals revealed a significant difference driven by “repeat-reward” (t-test: *t* = 7.68, *df* = 95, *p <* 0.0001), indicating that both explanations may be concomitant: a tendency to stick to rewarded choices, as well as a small but significant use of one-back stimulus and outcome information were used to guide choice. Like regression model 1, in regression model 2 the significant regressors in set size = 2, held for set size = 4, but this time only for male animals (Fig. S3B,E).

In regression model 3, we next looked at effects of correct history, reward, as well as repeat of previous reward and repeat of previous non-reward trials separately, reversing the role of reward and repeated cues used in the previous analysis. We found that all regression weights were positive (Fig. 3C,F). While the strong positive coefficient for reward in repeated trials from regression 2 seems to conform with reward-dependent repeating of a choice (with no regard to the cue), the positive effect for males of repeat from regression 3 in non-rewarded trials (Wilcoxon signed rank test, males: *W* = 1230, *p* = 0.02, t-test, females: *t* = 0.03, *df* = 38, *p* = 0.9) suggests that male mice might be able to leverage memory from the last trial even if trial *t−* 1 was not rewarded (i.e. to perform a “lose-switch” choice), confirming the conclusion of previous regression 2 that some form of adaptive one-back information is used. Similar results in set size = 4 were seen for both male and female animals, except for the positive effect for males of repeat in non-rewarded trials (Fig. S3C,F), possibly driven by a lower proportion of trials.

To investigate potential changes in an animal’s trial-by-trial behavior throughout its development, we employed mixed-effects models. In these models, each session’s regression coefficient served as the dependent variable, while session age was used as the predictor. This approach allowed us to examine how specific aspects of behavior, captured by the regression coefficient weights, varied across different developmental stages and enabled us to statistically account for repeated sessions by the same animal (see Mixed linear model analysis). Most significant developmental changes were seen in male, not female, animals (see Fig. 4,5). In the second regression for set size = 2, where trials were separated into repeat and non-repeat trials, the regression coefficient for repeated cues increased for males significantly over development (Fig. 4C, *β*_*age*_ = 0.024, 95 % CI = [0.016, 0.033], *p <* 0.0001). The contribution of repeat trials with a previous reward, also increased over development (Fig.4D: coefficient = 0.014, 95 % CI = [0.009, 0.018], *p <* 0.0001). This, combined with a decrease across development in the effect of reward in non-repeated cue trials for males (Fig. 4E: *β*_*age*_ = − 0.010, 95 % CI = [*−* 0.014, *−* 0.006], *p <* 0.0001), indicates that as male animals age, they are more likely to stick with their choice in trial t - 1 if they were rewarded, independently of the cue. In the third regression, while the main effect of reward was insignificant for both sexes (Fig. 4H, Fig. 5H), for males, the regressors that captured the repeating of a choice regardless of reward outcome, both increased over development (Fig. 4I: *β*_*age*_ = 0.03, 95% CI = [0.022, 0.038], *p <* 0.0001; Fig. 4J: *β*_*age*_ = 0.009, 95% CI = [0.004, 0.013], *p <* 0.0001). Interestingly, the influence of correct history did not change over development in either males or females, indicating that the contribution of RL-like learning in this simple associative learning task context is consistent across adolescence into early to adulthood.

### Computational Modeling

To build on our regression results and obtain a more quantitative and mechanistic understanding of potential one-back effects we next used computational modeling. The models we compared were generally based on RL tracking of stimulus-action values, but also included dependency on previous trials for choice policy (see Materials and methods). Using model comparison we identified a complex model with multiple parameterized one-back policies as the best fitting and most recoverable model (Fig. 6B). The winning model contained a single learning rate *α* and noise parameter *β*, with learning from negative outcomes *α*_*−*_ = 0; as well as three free parameters characterizing one-back policy bias as a function of last trial’s characteristics (see Fig. 6A for parameter schematic). These three one-back policy parameters emerged from an initial group of four parameters: two in response to non-rewarded outcomes (S1 and S3) and two in response to rewarded outcomes (S2 and S4). The first, parameter S1, or “Inappropriate Lose-Shift,” biased against repeating the previous trial’s choice when the stimulus changed and the animal’s previous choice was unrewarded. Parameter S3, or “Inappropriate Lose-Stay,” biased policy towards repeating the previous choice even when the stimulus stayed the same and the animal’s previous choice was unrewarded (repeating an incorrect choice), thus capturing perseveration. We initially considered separate S2 and S4 parameters (see Fig. 6A): “Inappropriate Win-Stay,” S2, captured staying with the previous trial’s choice after a rewarded trial even when the stimulus changed; and “Appropriate Win-Stay,” S4, indicated staying with the previous trial’s choice that was rewarded when the stimulus stayed the same. But, because S2 and S4 were highly correlated with each other, we reduced model complexity by setting S2=S4, a parameter we called “Stimulus Insensitive Win-Stay”. This parameter, biased towards repeating the previous trial’s choice after a win independently of whether the past stimulus changed or stayed the same.

We explored other models (including various parameterizations of last-trial-dependent policy biases), and found that this 5-parameter model was best able to recover the data (see Fig. 6E-K, Materials and methods, and Fig. S1). We also confirmed that model parameters were identifiable (See Materials and methods, Fig. S2); and that the model captured individual differences across mice and sessions accurately (Fig. 6G-K).

Once the data from each mouse and each session were fit using the winning model, we examined how the fit model parameters changed with age when mice were learning novel odor sets. The average learning rate *α*_+_ and softmax *β* parameter in set size = 2 and set size = 4 were stable across this adolescent period of development in both sexes (Fig. 7A-B,F-G; 8A-B,F-G) (mixed linear model, males: *α*_+_ parameter, set size = 2: *β*_*age*_ = *−*0.004, 95 % CI = [*−*0.025, 0.017], *p* = 0.716; *β* parameter, set size = 2: *β*_*age*_ = 0.002, 95% CI = [*−*0.006, 0.011], *p* = 0.637; males *α*_+_ parameter, set size = 4: *β*_*age*_ = *−*0.009, 95 % CI = [*−*0.033, 0.015], *p* = 0.450; *β* parameter, set size = 4: *β*_*age*_ = 0.007,95 % CI = [*−*0.005, 0.020], *p* = 0.251; females *α*_+_ parameter, set size = 2: *β*_*age*_ = *−*0.016, 95% CI = [*−*0.066, 0.035], *p* = 0.546; *β* parameter, set size = 2: *β*_*age*_ = 0.002, 95% CI = [*−*0.02, 0.02], *p* = 0.858, females *α*_+_ parameter, set size = 4: *β*_*age*_ = 0.012, 95 % CI = [*−*0.064, 0.088], *p* = 0.753; *β* parameter, set size = 4: *β*_*age*_ = 0.009, 95 % CI = [*−* 0.032, 0.050], *p* = 0.656).

**Fig 7.**
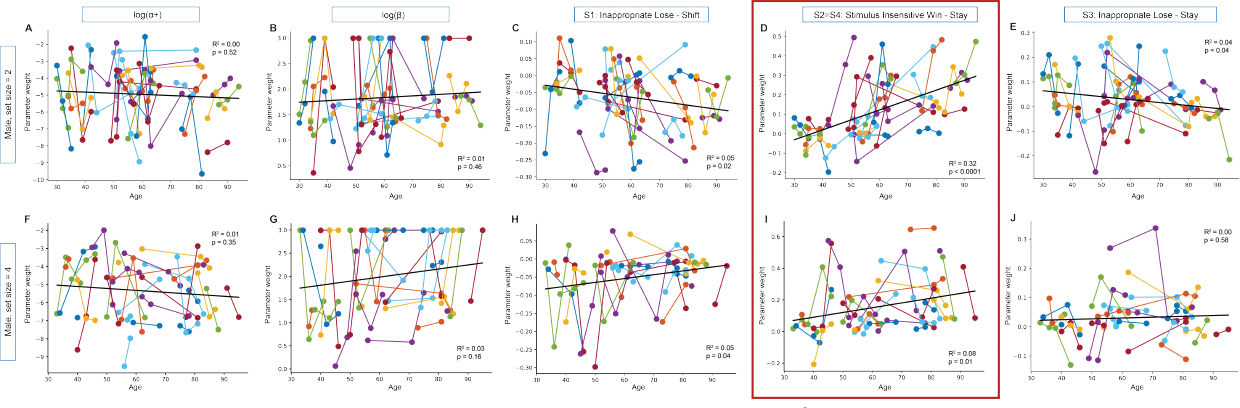
Strategy parameters across development for male mice in set size = 2 and set size = 4 from winning computational model. In order to understand the relationship between age and the parameters from our winning model (see Fig. 6 and Materials and methods for model details), we looked at whether parameter weight (y-axis) would change over development (x-axis) for male mice in set size = 2 (A-E) and set size = 4 (F-J). Parameter values are colored by individual with separate sessions connected by lines. For each parameter, we report the *R*^2^ and *p*-value from a simple linear regression (best fit line) and the results of a mixed linear regression model with age as the predictor variable and the parameter as the dependent variable in order to better account for variability driven by repeat sessions by individual mice (see Mixed linear model analysis for more information about the analyses). RL parameters such as *alpha*_+_ learning rate and softmax *beta* were stable across development in both set size = 2 (*alpha*_+_ (A): *β*_*age*_ = *−*0.004, 95 % CI = [*−*0.025, 0.017], *p* = 0.716; *β* (B): *β*_*age*_ = 0.002, 95% CI = [*−*0.006, 0.011], *p* = 0.637) and set size = 4 (*alpha*_+_ (F): *β*_*age*_ = *−* 0.009, 95 % CI = [*−* 0.033, 0.015], *p* = 0.450; *β* (G): *β*_*age*_ = 0.007,95 % CI = [*−* 0.005, 0.020], *p* = 0.251. Both parameters S1 “Inappropriate Lose-Shift” and S3 “Inappropriate Lose-Stay” decreased significantly with age in set size = 2 (S1 (C): *β*_*age*_ = *−*0.002, 95% CI = [*−*0.003, 0.0], *p* = 0.023; S3 (E): *β*_*age*_ = *−*0.001, 95% CI = [*−*0.003, 0.0], *p* = 0.04, but not in set size = 4 (S1 (H): *β*_*age*_ = 0.00, 95 % CI = [*−* 0.001, 0.002], *p* = 0.474; S3 (J): *β*_*age*_ = 0.00, 95 % CI = [*−* 0.002, 0.001], *p* = 0.607. However, strategy parameter, S2=S4 “Stimulus Insensitive Win Stay,” increased significantly for male mice in both set size = 2 (*β*_*age*_ = 0.006, 95% CI = [0.004, 0.007], *p <* 0.0001) and set size = 4 (*β*_*age*_ = 0.003, 95 % CI = [0.00, 0.006], *p* = 0.032).

The one-back strategy parameter that reflects perseveration after a rewarded trial, Parameter S2=S4: “Stimulus Insensitive Win-Stay,” grew with age in males in set size = 2 (mixed linear model, *β*_*age*_ = 0.006, 95% CI = [0.004, 0.007], *p <* 0.0001, Fig. 7D), and was replicated in set size = 4 (*β*_*age*_ = 0.003, 95 % CI = [0.00, 0.006], *p* = 0.032, Fig. 7I). Growth with age in the S2=S4 parameter was not seen in females in either set size (set size = 2:*β*_*age*_ = 0.001, 95% CI = [*−*0.004, 0.005], *p* = 0.713, Fig. 8D; set size = 4: *β*_*age*_ = 0.001, 95 % CI = [*−* 0.007, 0.010], *p* = 0.766, Fig. 8I).

**Fig 8.**
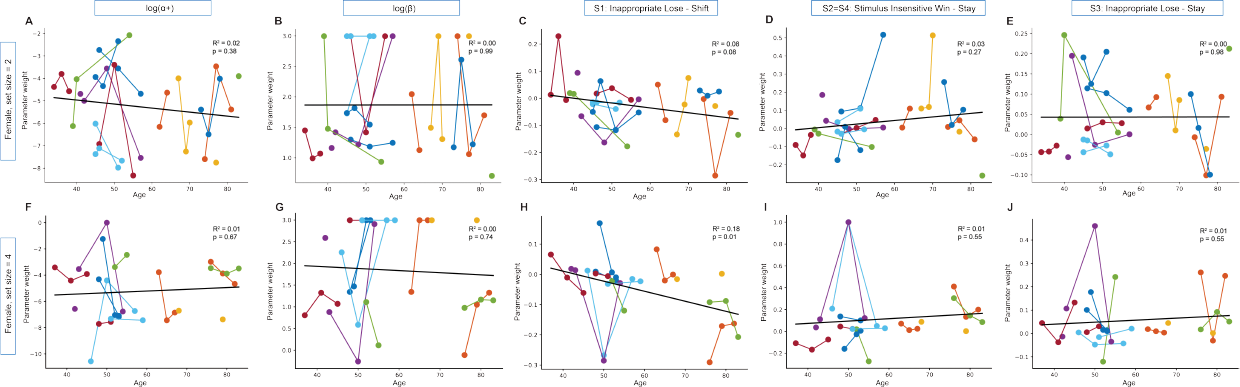
Strategy parameters across development for female mice in set size = 2 and set size = 4 from winning computational model. In order to understand the relationship between age and the parameters from our winning model (see Fig. 6 and Materials and methods for model details), we looked at whether parameter weight (y-axis) would change over development (x-axis) for female mice in set size = 2 (A-E) and set size = 4 (F-J). Parameter values are colored by individual with separate sessions connected by lines. For each parameter, we report the *R*^2^ and *p*-value from a simple linear regression (best fit line) and the results of a mixed linear regression model with age as the predictor variable and the parameter as the dependent variable in order to better account for variability driven by repeat sessions by individual mice (see Mixed linear model analysis for more information about the analyses). RL parameters such as *alpha*_+_ learning rate and softmax *beta* were stable across development in both set size = 2 (*alpha*_+_ (A): *β*_*age*_ = *−*0.016, 95% CI = [*−*0.066, 0.035], *p* = 0.546; *β* (B): *β*_*age*_ = 0.002, 95% CI = [*−*0.02, 0.02], *p* = 0.858) and set size = 4(*α*_+_ (F): *β*_*age*_ = 0.012, 95 % CI = [*−* 0.064, 0.088], *p* = 0.753; *β* (G): *β*_*age*_ = 0.009, 95 % CI = [*−* 0.032, 0.050], *p* = 0.656). Both parameters S2=S4 “Stimulus Insensitive Win Stay” and parameter S3 “Inappropriate Lose-Stay” did not change across development in either set size. (D) S2=S4, set size = 2: *β*_*age*_ = 0.001, 95% CI = [*−*0.004, 0.005], *p* = 0.713. (I) S2=S4, set size = 4: *β*_*age*_ = 0.001, 95 % CI = [*−*0.007, 0.010], *p* = 0.766. (E) S3, set size = 2: *β*_*age*_ = 0.00, 95% CI = [*−*0.003, 0.002], *p* = 0.74. (J) S3, set size = 4: *β*_*age*_ = 0.001, 95 % CI = [*−*0.002, 0.004], *p* = 0.556. However, female mice had a significant decrease in parameter S1 “Inappropriate Lose-Shift” in set size = 4 (H)(*β*_*age*_ = *−*0.003, 95 % CI = [*−*0.006, 0.00], *p* = 0.033) with a trend in a similar direction in set size = 2 (C)(*β*_*age*_ = *−*0.002, 95% CI = [*−*0.004, 0.00], *p* = 0.08).

The one-back strategy parameters S1, “Inappropriate Lose-Shift”, and S3,”Inappropriate Lose-Stay” indicate how an animal responds to a non-rewarded trial. Both of these parameters showed inconsistent age related changes between sexes and set sizes. Male animals in set size 2 had a significant decrease in age in parameter S1 (*β*_*age*_ = *−*0.002, 95% CI = [*−*0.003, 0.0], *p* = 0.023, Fig. 7C), but not in set size 4 (*β*_*age*_ = 0.00, 95 % CI = [*−*0.001, 0.002], *p* = 0.474, Fig. 7H). For parameter S3, male animals in set size 2 also had a significant decrease across development (*β*_*age*_ = *−*0.001, 95% CI = [*−* 0.003, 0.0], *p* = 0.04, Fig. 7E), but not in set size 4 (*β*_*age*_ = 0.00, 95 % CI = [*−* 0.002, 0.001], *p* = 0.607, Fig. 7J). We discounted these changes due to lack of consistency.

Female animals showed decreases in S1 with age that were at trend level in set size 2 and significant in set size 4(females S1, set size = 2: *β*_*age*_ = *−*0.002, 95% CI = [*−*0.004, 0.00], *p* = 0.08, Fig. 8C, females S1, set size = 4: *β*_*age*_ = *−*0.003, 95 % CI = [*−*0.006, 0.00], *p* = 0.033, Fig. 8H). In females the S3 parameter showed no change with age in either set size (females, S3, set size = 2: *β*_*age*_ = 0.00, 95% CI = [*−* 0.003, 0.002], *p* = 0.74, Fig. 8E, S3, set size = 4: *β*_*age*_ = 0.001, 95 % CI = [*−* 0.002, 0.004], *p* = 0.556, Fig. 8J).

We next looked for effects of session (meta-learning from repeat exposure) for each of the five parameters. Set size = 2 and set size = 4 sessions were interspersed so we combined and analyzed session data from set size = 2 and set size = 4 in the order mice experienced them. This revealed that alpha learning rate and beta did not show an effect of session for either sex (males, *alpha*_+_: *β*_*session*_ = *−*0.094, 95% CI = [*−*0.246, 0.058], *p* = 0.22; males, *beta*: *β*_*session*_ = 0.053, 95% CI = [*−*0.023, 0.128], *p* = 0.17; females, *alpha*_+_: *β*_*session*_ = *−*0.022, 95% CI = [*−* 0.304, 0.261], *p* = 0.88; females, *beta*: *β*_*session*_ = 0.040, 95% CI = [*−* 0.089, 0.168], *p* = 0.54, Fig. 9A-B,F-G). However, use of the three one-back parameters changed significantly across sessions in males with S1 “Inappropriate Lose Shift” decreasing across sessions (*β*_*session*_ = *−*0.010, 95% CI = [*−*0.018, *−*0.002], *p* = 0.01), S2 = S4 “Stimulus Insensitive Win Stay” increasing (*β*_*session*_ = 0.018, 95% CI = [0.005, 0.032], *p* = 0.009), and S3 “Inappropriate Lose Stay” decreasing (*β*_*session*_ =*−* 0.018, 95% CI = [*−* 0.027, *−* 0.009], *p <* 0.0001). As could be expected, these experience-dependent trends are in the same direction as male set size = 2 changes over age (Fig. 7C-E), although this trend is less captured by set size = 2 (Fig. 7H-J). Together, the opposing trends in S1 and S2 = S4 could indicate that male mice are more likely to stick with their previous choice, regardless of reward or stimuli, as they gain experience in the task (see 6A). A decrease in S3 across sessions shows that males also switch their response following an unrewarded trial when the stimuli stays the same, which may reveal an increase in task knowledge over time. Females, showed a trend toward a decrease in S1 parameter weights over sessions, similar to males, but no significant changes across sessions in one-back strategies (S1: *β*_*session*_ = *−*0.012, 95% CI = [*−*0.026, 0.001], *p* = 0.068; S2=S4: *β*_*session*_ = 0.013, 95% CI = [*−* 0.019, 0.044], *p* = 0.42; S3: *β*_*session*_ = − 0.003, 95% CI = [*−* 0.018, 0.013], *p* = 0.75, Fig.9H-J). This is similar to small, if any, changes across age found in female mice (Fig. 8C-E;H-J).

**Fig 9.**
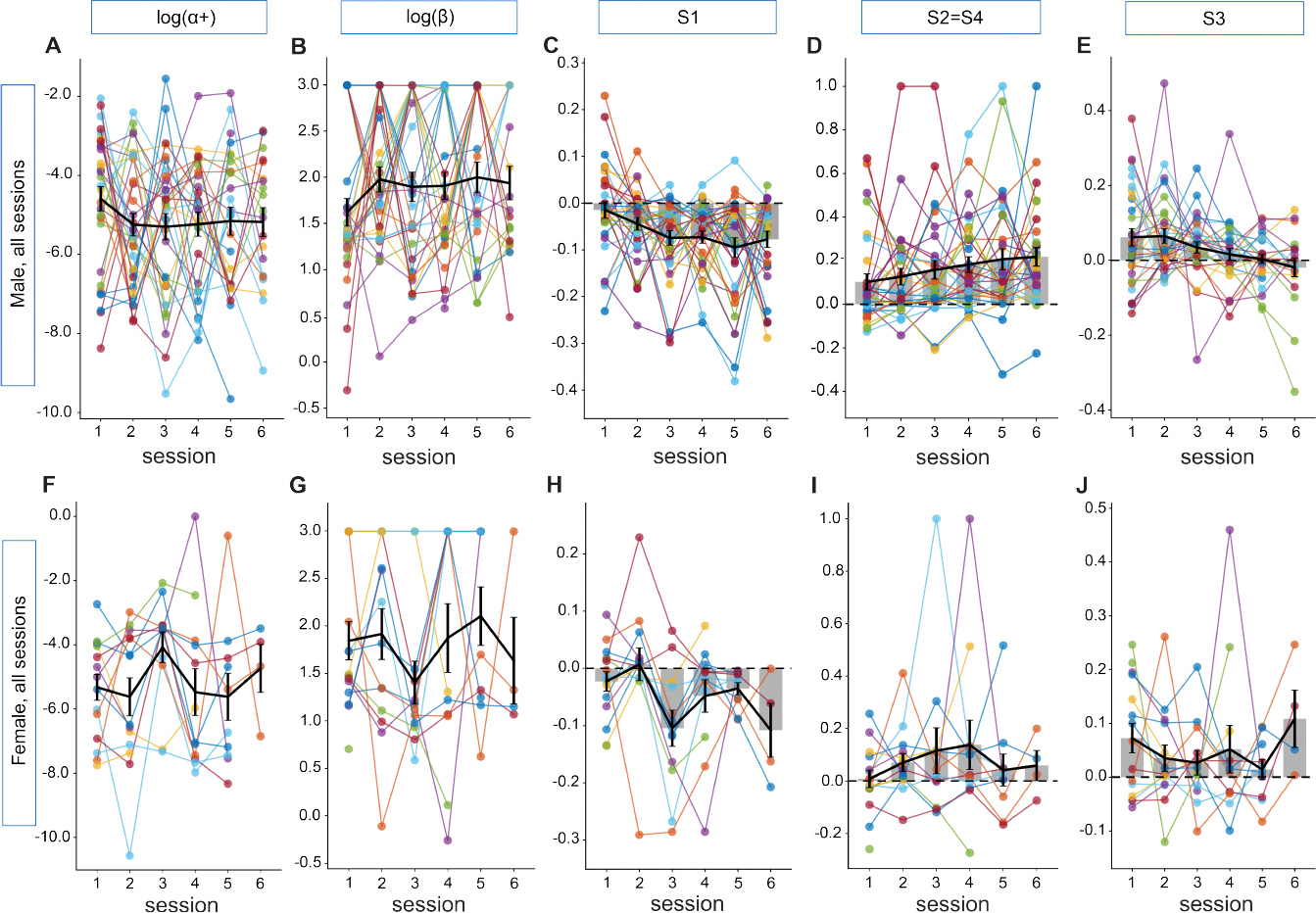
Effect of session on winning model parameters for set size = 2 and set size = 4 for both male and females. To test if mice adjusted one-back strategies with experience, we next compared how parameter weights (y-axis) changed across the 6 sessions analyzed for each mouse (x-axis). Since set size = 2 and set size = 4 days were interspersed, we combined set size data and analyzed sessions chronologically. Since each mouse had multiple sessions, each individual was colored and their sessions connected, and mixed linear models were used to account for individual variability (see Mixed linear model analysis for more information about analysis). Gray bars indicate mean values for each session with SEM and a line connecting each mean. Dotted lines are set to 0 for strategy parameters. Similar to what was seen in comparisons between regression coefficients and age, there was no effect of experience on *alpha*_+_ or *beta* parameters for either sex. For male mice (A) *alpha*_+_: *β*_*session*_ = *−*0.094, 95% CI = [*−*0.246, 0.058], *p* = 0.22; (B) *beta*: *β*_*session*_ = 0.053, 95% CI = [*−*0.023, 0.128], *p* = 0.17. For female mice (F) *alpha*_+_: *β*_*session*_ = *−*0.022, 95% CI = [*−* 0.304, 0.261], *p* = 0.88; (G) *beta*: *β*_*session*_ = 0.040, 95% CI = [*−* 0.089, 0.168], *p* = 0.54. Use of one-back strategy parameters changed significantly across sessions for male mice with (C) S1 “Inappropriate Lose Shift” decreasing across sessions (*β*_*session*_ = *−*0.010, 95% CI = [*−*0.018, *−*0.002], *p* = 0.01), (D) S2 = S4 “Stimulus Insensitive Win Stay” increasing (*β*_*session*_ = 0.018, 95% CI = [0.005, 0.032], *p* = 0.009), and (E) S3 “Inappropriate Lose Stay” decreasing (*β*_*session*_ = *−* 0.018, 95% CI = [*−* 0.027, 0.009], *p <* 0.0001). Female mice showed a trend in the same direction of male mice in (H) S1 “Inappropriate Lose Shift” (*β*_*session*_ = *−* 0.012, 95% CI = [*−* 0.026, 0.001], *p* = 0.068), but did not show any experience related changes in (I) S2=S4 “Stimulus Insensitive Win Stay” (*β*_*session*_ = 0.013, 95% CI = [*−*0.019, 0.044], *p* = 0.42) or (J) S3 “Inappropriate Lose Stay” (*β*_*session*_ = *−*0.003, 95% CI = [*−*0.018, 0.013], *p* = 0.75).

These modeling results suggest that mice have access to, and leverage information from, short-term memory for making choices in learning situations, as humans do. However, they use this information for short-term one-back choice strategies instead of human-like working memory.

## Discussion

We tested mice in a conditional associative learning task with a set size manipulation that has been used previously to disentangle contributions of working memory (WM) from reinforcement learning (RL) systems. A goal of our study was to identify the extent to which mice use WM and the age at which both WM and RL become adult-like.

Mice readily learned the odor-based task when presented with both set size = 2 and set size = 4 sessions, moving gradually from chance performance to above 70% correct over the course of 200 trials per odor. While in the last 50 trials per odor male animals performed significantly better in set size = 2 than set size = 4 (Fig. 2A), this was likely driven by more repeat trials in set size = 2 than set size =4, rather than a notable learning differences between set sizes. Gradual learning of stimulus-action associations is consistent with the use of an RL strategy ([10]) and this, combined with the learning similarity between set sizes, was an initial clue that mice may not be using canonical WM to solve the task. However, we also noted better performance for male animals in trials when an odor was repeated over non-repeated trials across both set sizes (Fig. 2B), a difference which could either be evidence of one-trial-back WM or consistent with a non-WM dependent one-trial-back strategy. To parse these differences in strategy, we performed regression analyses and fit trial-by-trial RL models.

Our regression analyses confirmed that history of rewarded trials, a marker of RL, and certain patterns of repeated, rewarded, and both non-repeated and non-rewarded trials contributed significantly to performance (see Figs. 3, S3). Next, by using RL models, we found that along with more standard parameters like learning rate *α* and softmax *β*, the inclusion of parameters weighing a variety of one-back strategies allowed better fit and recovery of behavioral data for individual mice. Together, these data suggest mouse performance in the RL+WM task can be well explained by RL and simple one-back choice strategies.

While our data show that mice use one-trial-back information to influence their decision above and beyond cumulative reward, this does not appear to benefit their performance in this task, suggesting that mice may not rely on canonical working memory in the way observed in human learning ([8]). This is surprising in light of other studies that examine spontaneous alternation or choices informed by information that has to be maintained over a delay in rodents ([6, 7]). Successful performance on these tasks suggest there is some form of working memory available to rodents that is not recruited in our task. This could be driven, in part, by our task design. For example, two or four novel odors are paired daily with only two lateral ports, allowing mice to use a multitude of strategies instead of forcing a choice as in spontaneous alternation tasks. In addition, the presentation of multiple stimuli from a central port in an animal-directed and semi-random manner allowed us to parse spatial strategies (such as orienting behavior) from stimulus-dependent behavior, reducing the likelihood of conflating non-WM strategies with WM ([4]).

Because we tested mice across adolescent to adult ages, we were able to examine the effect of age on task and model metrics. We found a consistently significant and positive age-related relationship in male animals in the S2=S4: Stimulus Insensitive Win-Stay. This indicates that male mice become more likely to stay following a reward as they develop. Increases in use of S2=S4 win-stay strategy was not seen with age in female mice. We speculate this sex difference may result from greater energy costs for moving in males due to their heavier weights and/or steeper growth curves ([15].

We also found evidence that mice relied on one-back strategies after non-rewarded trials (S1 and S3). Use of these strategies showed complex patterns depending on sex and age that were not fully consistent across both set sizes. Use of these strategies also changed significantly with session experience in males with a similar trend in females showing a decrease in S1,”Inappropriate Lose-Shift.” This decrease in S1 could be interpreted as growing knowledge of the task and, in conjunction with an increase in parameter S2 = S4, could indicate a increase in staying with the previous choice, regardless of a previous stimuli or reward.

Our data did not reveal any significant developmental changes in basic performance metrics, learning rate *α*, and the softmax inverse temperature parameter *β*, with age. The lack of change in these metrics across adolescence was unexpected and may indicate mouse-human differences. In many tasks across a range of species adolescent changes in learning and working memory are well established ([16]; [17]; [18]; [19]). For example, in rats, there is improvement in a delayed match to sample task across adolescence and into adulthood with time delays of up to 24 seconds ([6]). In non-human primates, adult animals had higher performance than adolescents in an oculomotor delayed response task, a task thought to measure working memory ([17]). Similarly, not only working memory, but reinforcement learning metrics such as, *β* and *α* learning rate, have been shown to shift across development in humans (reviewed [20]; see also [21]). The fact that our data shows little change in mice over development could be due to an emergence of adult-like function earlier in life to support dispersal and the transition to independence ([19]). It is also possible that mice use less working memory than rats due to a relatively less developed prefrontal cortex ([22]; [23]), the brain region known across species to be involved in working memory. Future work using the RL+WM task in rats will be needed to test if rats use RL plus WM or more one shot strategies in decision policies to navigate this simple learning task.

One design aspect of our experiment that may be an important caveat for comparisons to others’ data is the ‘readiness check.’ We started each daily session by having mice perform the task with odors A & B, only moving them on to learning novel odors when they passed a performance criterion (see Materials and methods). While this allowed us to ensure we were studying novel odor learning in mice on days when they had strong motivation, it may filter out participation on days with a less efficient state for learning, effectively equalizing a baseline state. This kind of readiness check is not often included in other learning tasks for rodents and therefore needs to be considered when comparing data across tasks and labs as well as species.

Our research highlights the importance of considering multiple components contributing to learning behavior in rodents. Recent research has shown evidence that animals do not only rely on single strategies, but use a wide repertoire during decision making ([24]). Here, we also consider how multiple strategies contribute to decision making in the context of learning, but instead of assuming that animals switch between them over the course of the experiment, we show how they are used concomitantly throughout a learning session. Thus our approach differs from others ([25], [24], [26]) by looking at how multiple parallel processes’ contribute to behavior, rather than multiple strategies supported by the same process.

## Conclusion

In conclusion, our study establishes male and female mice at different stages of development use similar strategies to learn a conditional associative task with multiple stimuli and two alternative choices. In depth modeling of the data suggest mice do use reinforcement learning but also rely on one-back strategies to solve the task.

## Supporting information

**Fig S1.**
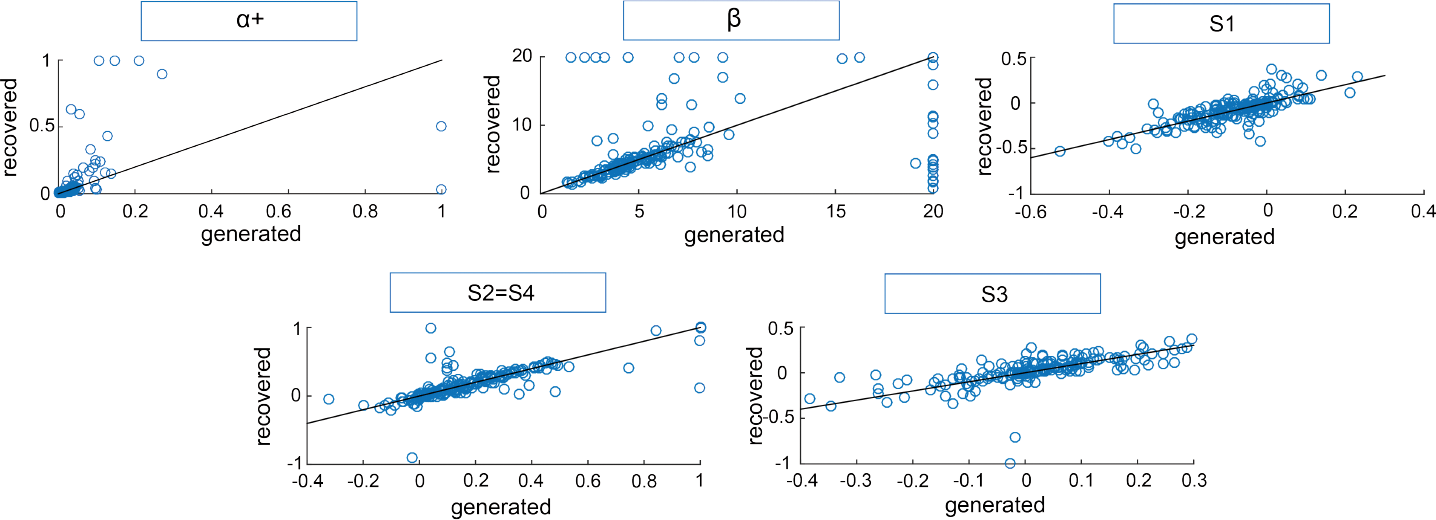
The winning model, a0b1232, parameters are identifiable. We tested a series of RL-based computational models that explored which strategies mice may use to integrate and use past trial information for trial-by-trial learning. The winning model, as shown by BIC and protected exceedence probability (see Fig.6B,C), had 5 free parameters that included *alpha*_+_ (a), no *alpha*_*−*_ (0), a softmax *beta* parameter (b) and 3 one-back strategy parameters (see Fig. 6A for schematic, Materials and methods for more details). In order to validate our winning model, we generated data by simulating the model with parameters fit on inidivdual sessions. Then, we fit the simulated data in order to obtain recovered parameters. Recovered model parameters (y-axis) were highly correlated with generating parameters (x-axis), indicating that model parameters are identifiable ([12]).

**Fig S2.**
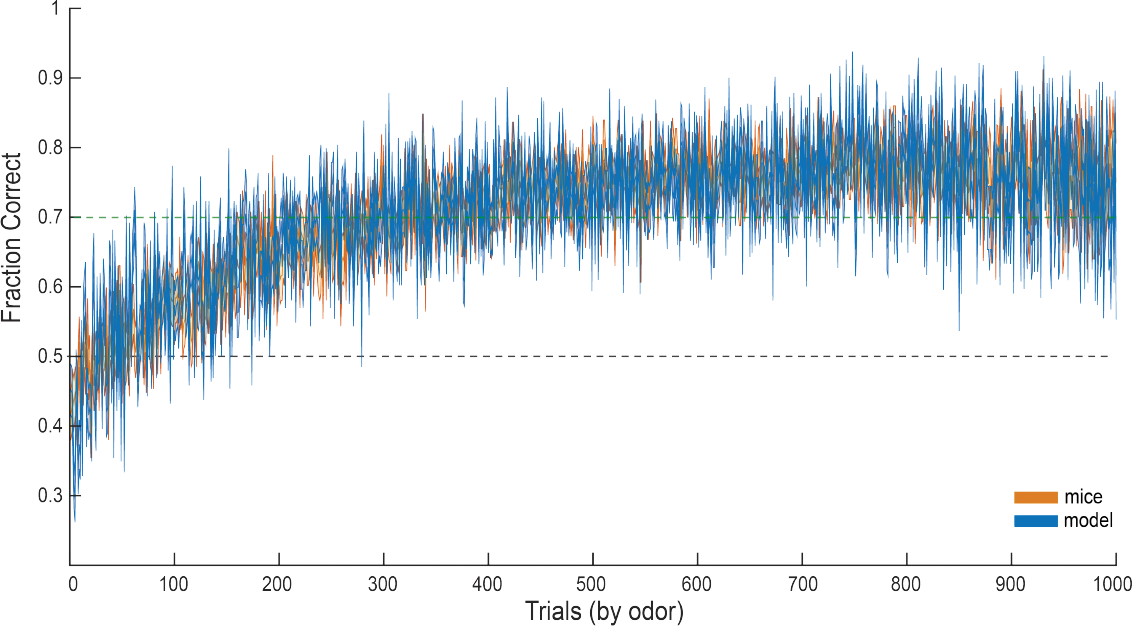
The winning model, a0b1232, captures group learning. In order for a model to be valid, it is important that it captures both trial-by-trial learning as well as learning that takes place across a session. Our winning model, a0b1232, includes an RL-learning component as described in Materials and methods along with 5 free parameters as described in the name. *alpha*_+_ is indicated by (a), softmax *beta* parameter by (b), the absence of any *alpha*_*−*_ learning rate is indicated by (0). The final model also included 3 one-back strategy parameters (S1, S2=S4, S3, see Fig. 6A for schematic) that captured how mice’ responses to a current trial reflect one-back stimuli and reward experience. Here, we generated data by simulating the model with parameters fit on individual sessions. Then, we inspected the similarity between simulated data (blue) and mouse data (orange) and found that the model simulated with fit model parameters captures well mouse learning curve data.

**Fig S3.**
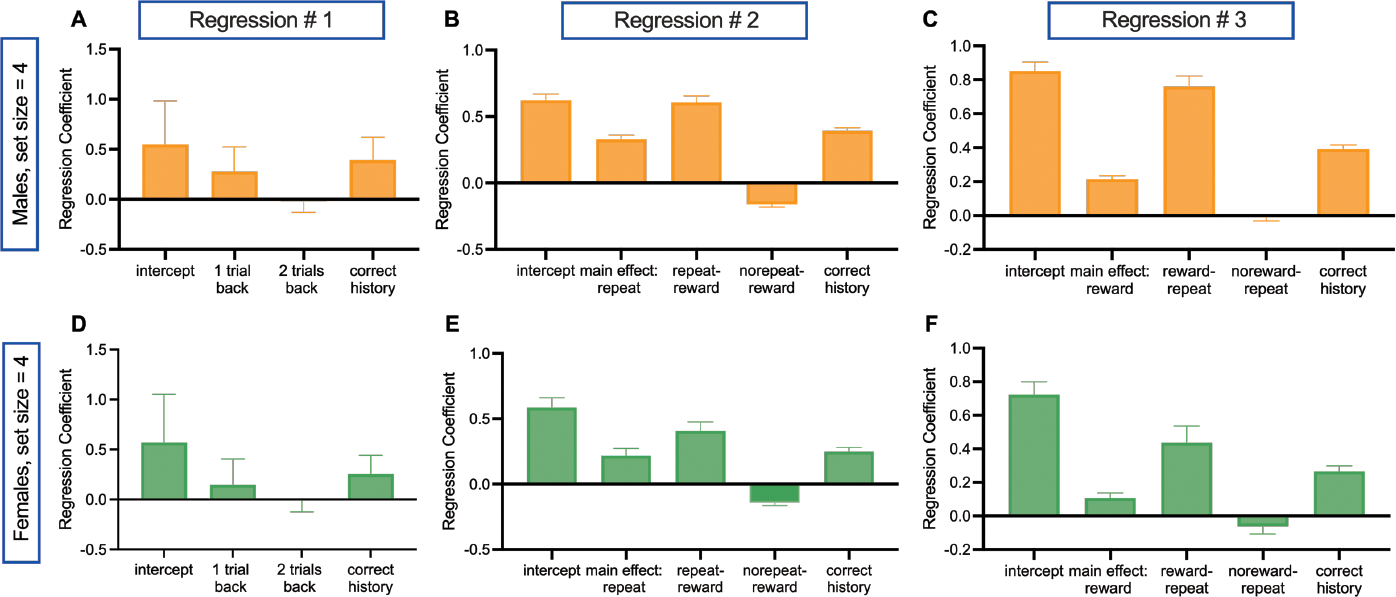
Summary of logistic regression coefficients for males and females in set size = 4. We used logistic regression analyses to better characterize trial-by-trial dynamics in behavior. We developed three different logistic regressions to capture differing hypotheses about which trials and what kinds of trials drive choice behavior on trial *t*. All regressions used correct history, a theoretical marker for incremental RL, and a variety of predictors that rely on more recent information which may underlie WM. ‘Correct history’ was a cumulative sum of all past correct choices for the current stimuli on trial *t*. Our first logistic regression for males (A) and females (D) looked at the influence of correct history and either one or two trials back on correct choice. For males: correct history: *t* = 15.13, *df* = 77, *p <* 0.0001; 1-trial back: *t* = 10.62, *df* = 77, *p <* 0.0001; 2-trials back: *t* = 1.631, *df* = 77, *p* = 0.10. And for females: correct history: *W* = 358.0, *p <* 0.0001; 1-trial back: *W* = 248.0, *p* = 0.002; 2-trials back: *W* = *−* 28, *p* = 0.75. The second logistic regression for males (B) and females (E) kept ‘correct history’ as a predictor, but now differentiated one-back trials into repeat stimuli, using a main effect and two interaction terms (repeat - reward and no repeat - reward) to see whether stimulus identity could predict choice. For males: correct history: *t* = 14.88, *df* = 77, *p <* 0.0001; main effect of repeat: *t* = 10.60, *df* = 77, *p <* 0.0001; repeat - reward: *t* = 11.24, *df* = 77, *p <* 0.0001; no repeat - reward: *t* = 6.94, *df* = 77, *p <* 0.0001. And for females: correct history: *t* = 7.0, *df* = 26, *p <* 0.0001; main effect of repeat: *W* = 248, *p* = 0.002; repeat - reward: *t* = 4.826, *df* = 26, *p <* 0.0001; no repeat - reward: *t* = 4.52, *df* = 26, *p* = 0.0001. For our final logistic regression in males (C) and females (F) we kept ‘correct history’ and re-organized the trials to reflect whether or not a mouse was rewarded for their choice in one-back trials (instead of looking at the stimulus) by using a main effect and two interaction (reward - repeat and no reward - repeat). For males: correct history: *t* = 14.88, *df* = 77, *p <* 0.0001; main effect of reward: *W* = 2721, *p <* 0.0001; reward - repeat: *t* = 11.76, *df* = 77, *p <* 0.0001; no reward - repeat: *t* = 0.18, *df* = 77, *p* = 0.85. For females: correct history: *t* = 7.0, *df* = 26, *p <* 0.0001; main effect of reward: *t* = 3.93, *df* = 26, *p* = 0.0006; reward - repeat: *t* = 4.34, *df* = 26, *p* = 0.0002, no reward - repeat: *t* = 0.90, *df* = 26, *p* = 0.37. Together these results indicate a significant influence of both correct history (RL) and combinations of one-back trials on current mouse choice. Note that for our second regression for set size = 4, the absolute number of repeat trials are fewer than set size = 2.

**Fig S4.**
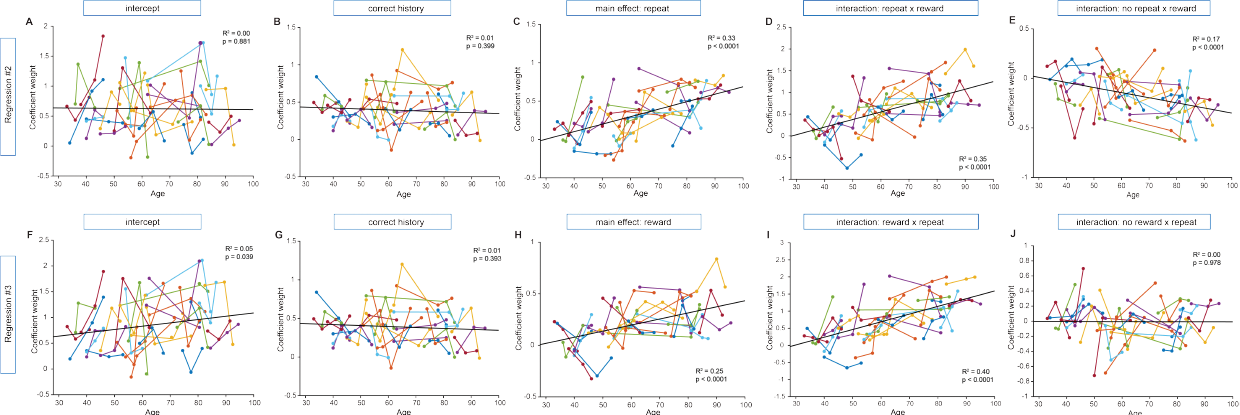
Male regression coefficients across development in set size = 4. In order to understand the relationship between age and the predictors of current choice in trial *t* (see Fig. S3 for the logistic regression summary), we looked at whether coefficient weight (y-axis) would change over development (x-axis) for male mice. It is possible that as mice age, they rely to differing extents on separable learning systems (for example, RL, captured by ‘correct history’ and, WM, captured by one-back combinations of stimuli and reward). Here we report the *R*^2^ and *p*-value from a simple linear regression (best fit line) on each figure and the results of a mixed linear regression model that better allowed us to account for any variability caused by repeat sessions of individual mice (repeat sessions are connected with a line; see Mixed linear model analysis for more information about analyses). Our second regression (A-E), focused on the influence of repeating or non-repeating stimuli in one-back trials on current choice on repeat trials, (A) intercept: *β*_*age*_ = 0.00, 95 % CI = [*−* 0.006, 0.006], *p* = 0.994, while our third regression (F-J), focused on the influence of one-back rewarded or non-rewarded trials influenced the mouse’s current choice, (F) intercept: *β*_*age*_ = 0.008, 95% CI = [0.001, 0.016], *p* = 0.024. Despite ‘correct history’ being a significant predictor of current choice in both regression #2 and regression #3 (S3B,C), ‘correct history’ did not change over development: (B) regression #2, correct history: *β*_*age*_ = 0.002, 95%CI = [*−* 0.004, 0.008], *p* = 0.561, and (G) regression #3, correct history: *β*_*age*_ = *−* 0.001, 95% CI = [*−* 0.004, 0.002], *p* = 0.509. The main effect regression weight of both reward and whether the stimuli repeated on trial *t* increased significantly over time: (C) regression #2, main effect of repeat: *β*_*age*_ = 0.008, 95% CI = [0.005, 0.011], *p <* 0.0001, and (H) regression #3, main effect of reward: *β*_*age*_ = 0.005, 95% CI = [0.003, 0.008], *p <* 0.0001 and interactions between repeat stimuli and previous reward (D: repeat-reward for regression #2 and I: reward - repeat for regression #3) also increased significantly over time: (D) repeat - reward: *β*_*age*_ = 0.011, 95% CI = [0.004, 0.019], *p* = 0.004; (I) reward - repeat: *β*_*age*_ = 0.020, 95% CI = [0.013, 0.028], *p <* 0.0001. The increase over development in the regression weights of both main effect and repeat-reward interactions could indicate that some form of adaptive one-back strategies increases across development. This, combined with the significant negative direction of no repeat - reward (E: *β*_*age*_ = *−* 0.005, 95%CI = [*−* 0.008, *−* 0.002], *p* = 0.001) could indicate that as male mice age they are more likely to stick to their previous choice. Interestingly, unlike in set size = 2 (4J), there was no significant increase in the regression weight for no reward - repeat (J: *β*_*age*_ = 0.00, 95% CI = [*−* 0.004, 0.003], *p* = 0.818) which could be influenced by the addition of two more stimuli. With this single exception, all developmental changes seen in set size = 4 were also reflected in set size = 2 4.

**Fig S5.**
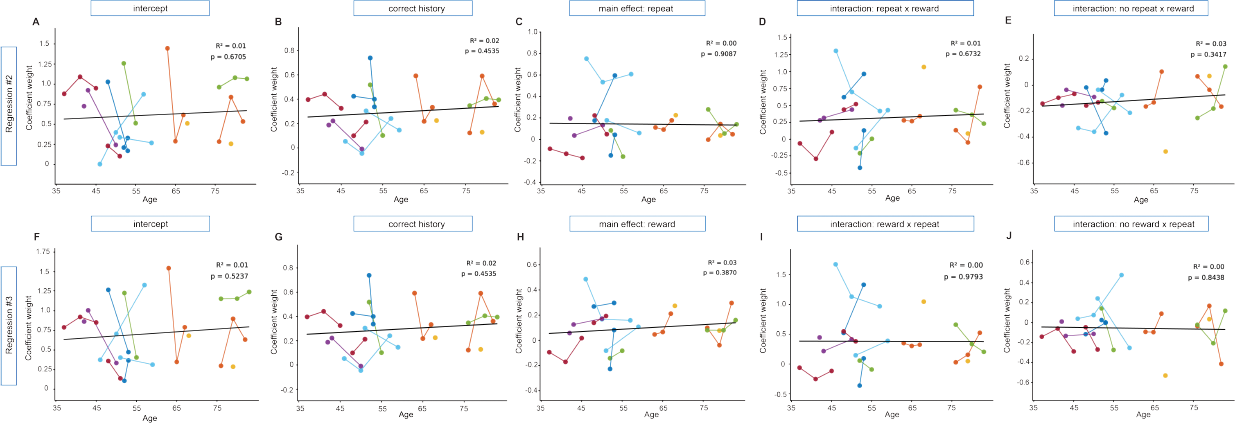
Female regression coefficients across development in set size = 4. Multiple logistic regressions were used to determine the influence of recent information on trial *t*, a marker for WM-like learning, and the influence of past rewarded trials for the same stimulus at trial *t*, a marker for RL (Fig. S3). In order to determine which, if any, regression coefficients changed across development for female mice in set size = 4, we looked at the relationship between age at session (x-axis) and coefficient weight (y-axis). Here, similar to Fig. S4 we report the *R*^2^ and *p*-value from a simple linear regression (best fit line) on each figure and the results of a mixed linear regression model that better allowed us to account for any variability caused by repeat sessions of individual mice (repeat sessions are connected with a line; see Mixed linear model analysis for more information about analyses). Our second regression (A-E), focused on the influence of repeating or non-repeating stimuli in one-back trials on current choice on repeat trials, (A) intercept: *β*_*age*_ = 0.002, 95 % CI = [*−* 0.001, 0.013], *p* = 0.797, while our third regression (F-J), focused on the influence of one-back rewarded or non-rewarded trials influenced the mouse’s current choice, (F) intercept: *β*_*age*_ = 0.003, 95 % CI = [*−* 0.009, 0.015], *p* = 0.628. Like male mice in both regression #2 and regression #3, the regression coefficient that is a marker for RL, “correct history,” was significantly above zero (see Fig. S3E,F). However, correct history, was steady across development in both regressions (regression #2 (B): *β*_*age*_ = 0.001,95 % CI = [*−* 0.004, 0.007], *p* = 0.605; regression #3 (G): *β*_*age*_ = 0.001, 95 % CI = [*−* 0.004, 0.007], *p* = 0.605). Unlike male mice (see S4), however, there was no significant changes in regression coefficient weight for female mice across development for any main effect or interaction. For regression #2, a regression focusing on the influence of stimuli on trial *t*: (C) main effect of repeat: *β*_*age*_ = *−*0.001, 95 % CI = [*−*0.009, 0.006], *p* = 0.735 (D) repeat - reward: *β*_*age*_ = 0.003, 95 % CI = [*−*0.011, 0.016], *p* = 0.683 (E) no repeat - reward: *β*_*age*_ = 0.002, 95 % CI = [*−*0.002, 0.006], *p* = 0.337. Similarly, for regression #3, a regression focusing on the influence of reward on trial *t*: H) main effect of reward: *β*_*age*_ = 0.002, 95 % CI = [*−*0.003, 0.007], *p* = 0.42, (I) reward - repeat: *β*_*age*_ = *−*0.001, 95 % CI = [*−* 0.017, 0.015], *p* = 0.919, (J) no reward - repeat: *β*_*age*_ = *−* 0.001, 95 % CI = [*−* 0.006, 0.005], *p* = 0.843. While it is possible that a low number of set size = 4 sessions for female mice may be responsible for no trial-by-trial learning changes over development, our findings replicate set size = 2 sessions for female mice (Fig. 5).

## Acknowledgments

This work was funded by NSF SL-CN: Science of Learning in Adolescence: Integrating Developmental Studies in Animals and Humans (Award #1640885 to L.W. and A.G.E.C.) and the Simons SFARI Foundation (award #613972 to L.W.). These foundations had no role in study design, data collection and analysis, decision to publish, or preparation of the manuscript.

